# Low-temperature inkjet-printed electrochemical sensors on OSTE+ microfluidics for oxygen monitoring and scavenging

**DOI:** 10.1101/2025.09.23.675348

**Authors:** Denise Marrero, Ferran Pujol-Vila, Eva Tuset, Gemma Gabriel, Rosa Villa, Mar Alvarez, Xavi Illa

## Abstract

Real-time monitoring of biological events is essential for advancing in drug testing, disease modeling, and precision medicine. However, conventional analytical methods often lack the spatiotemporal resolution to capture dynamic cellular responses, particularly in Organ-on-a-Chip (OoC) platforms and microphysiological systems (MPS). While progress has been made in sensor development, integrating sensors into microfluidic devices remains technically challenging. Materials commonly used in microfluidics, such as polydimethylsiloxane (PDMS), are poorly suited for stable sensor integration due to low metal adhesion. Off-stoichiometry thiol-ene-epoxy (OSTE+) polymers present a promising alternative due to the presence of unreacted thiol groups. These groups enable dual functionalities: strong metal affinity for direct for robust sensor integration, and oxygen scavenging essential for microscale oxygen level control. However, their compatibility with conventional sensor fabrication methods, particularly those requiring elevated temperatures, is limited. This work introduces a low-temperature fabrication method based on inkjet printing and photonic curing, suitable for thermally sensitive substrates. Gold and silver nanoparticle inks were directly sintered onto OSTE+ using photonic curing, producing functional electrochemical sensors for spatially resolved and real-time oxygen detection. Unlike conventional thermal sintering, which is incompatible with many microfluidic materials, photonic curing preserves the chemical reactivity and mechanical properties of the polymer without requiring surface treatment or adhesive layers. To demonstrate functionality, sensors were integrated into microfluidic devices for continuous monitoring of dissolved oxygen. Real-time oxygen depletion driven by the material’s intrinsic scavenging capability was successfully quantified. These findings support the compatibility of sensor integration into OSTE+ polymers enhancing the real-time monitoring capabilities in OoC and MPS systems.

**Highlights:**

- Inkjet-printed sensors are integrated into OSTE+ using a low-temperature protocol.
- Photonic curing enables sintering of gold and silver nanoparticle inks without damaging the polymer substrate.
- Oxygen scavenging is monitored in real time using integrated electrochemical sensors.
- Tunable oxygen gradients are achieved in OSTE+ microfluidics for organ-on-chip models.

## Introduction

The integration of sensors into microfluidic systems, and particularly microphysiological systems (MPS) and Organ-on-a-Chip (OoC) platforms, is essential for real-time, on-chip monitoring of dynamic biological events [1,2]. These events are often difficult to assess using conventional techniques, including microscopy, biochemical assays, and culture-based methods [3,4]. Such approaches provide valuable end-point or bulk measurements, but they often require off-chip processing that delays decision-making, and some rely on experimental termination, which prevents capturing transient responses and limits spatiotemporal resolution. Despite significant advances in sensor technologies, including electrical [5–7], electrochemical [8–10], optical [11–13], and mechanical sensors [14,15], their integration into microfluidic platforms remains challenging. Key challenges include poor adhesion of conductive materials to polymeric substrates, incompatibility of fabrication methods with thermally or chemically sensitive materials, and sensor delamination during long-term culture. Existing approaches such as bonding commercial sensors to chambers or introducing wires through device walls [16–18], compromise device miniaturization, limit multiplexing, and often introduce leakage or contamination risks. Addressing these limitations is critical to fully realizing the potential of MPS and OoC platforms for applications in drug testing, disease modeling and precision medicine.

Among the materials commonly used in microfluidic devices, polydimethylsiloxane (PDMS) is still the gold standard polymer for device fabrication due to its biocompatibility, optical transparency, flexibility and ease of processing via soft lithography. Despite its advantages, it presents several drawbacks both for biological modeling and sensor integration. PDMS is prone to absorbing small molecules, such as drugs, which can alter their bioavailability and compromise assay accuracy [19]. Additionally, its low surface energy and chemically inert methyl groups result in poor adhesion to metals, posing challenges for robust and stable sensor integration[20]. To address these limitations, hybrid fabrication strategies often incorporate materials with better adhesion to metals for electrode integration into the substrate while maintaining PDMS for other structural components. Metal patterning on PDMS remains challenging due to poor adhesion, often necessitating surface modifications such as oxygen plasma or the deposition of adhesion-promoting layers [21,22]. However, these approaches typically provide only temporary adhesion due to surface hydrophobic recovery and mechanical stress-induced delamination [23]. This limitation underscores the need for alternative materials that support both stable sensor integration and compatibility with cell culture environments.

Among these alternatives, off-stoichiometry thiol-ene-epoxy (OSTE+) polymers have emerged as a promising substitute for PDMS [24,25]. OSTE+ polymers are thermosetting materials that can also be fabricated by soft lithography. They are synthesized from allyl, thiol, and epoxy monomers in an off-stoichiometric ratio, and are cross-linked via UV-initiated click chemistry. A key advantage of OSTE+ formulations lies in their tunable two-step fabrication process. After the initial UV-induced polymerization, a partially cured network is formed, leaving reactive groups (such as thiols or epoxies) available for post-processing steps such as functionalization and bonding [26–28]. This is followed by a thermal curing step which completes the crosslinking, resulting in a chemically inert material. Unlike PDMS, OSTE+ polymers exhibit minimal absorption of small molecules, which is advantageous for applications involving drug transport studies [29]. Additionally, the strong affinity of thiol groups for noble metals, such as gold and silver, makes OSTE+ polymers a promising substrate for sensor integration, supporting stable monitoring in MPS and OoC systems. [30,31]. Moreover, OSTE+ polymers possess intrinsic oxygen scavenging properties, which can be used to regulate oxygen levels in MPS and OoC devices, offering a potential advantage in hypoxia and physiological microenvironment modeling [32,33]. This property originates from the presence of unreacted thiol groups at the surface and within the bulk material [34]. This oxygen scavenging capability was first explored by Sticker and colleagues in 2019 for cell culture applications, where the temperature of the thermal cure was adjusted to tune the oxygen scavenging efficiency [35]. Notably, the oxygen scavenging efficiency is inversely correlated with curing temperature ranging from 110°C to room temperature, with lower temperatures enhancing this property.

Despite the promising potential of this material family for sensor integration, developing a room-temperature fabrication strategy to maximize the oxygen scavenging properties of OSTE+ polymers while avoiding harsh chemicals and mechanical stress remains challenging. In this context, rapid prototyping techniques, such as the non-contact inkjet printing, offers a viable solution. Inkjet printing enables precise deposition of functional inks without the need for masks, facilitating the rapid prototyping of sensors under gentle conditions while minimizing material waste. However, a key limitation is its reliance on thermal sintering, typically performed at temperatures around 120-150°C [10]. This restricts the choice of materials to those with a glass transition temperature above the sintering temperature, rendering it incompatible with OSTE+ polymers. To address these limitations, photonic sintering has emerged as a promising alternative. This technique uses high-energy flash lamps to deliver intense, localized light pulses that rapidly sinter thin films without exposing temperature-sensitive substrates to the high temperatures of conventional thermal processes[36]. By enabling solvent evaporation and metal particle sintering in milliseconds, photonic curing minimizes thermal stress, making it an ideal curing strategy for integrating sensors into temperature-sensitive materials.

This work presents a low-temperature fabrication method that combines inkjet printing with photonic curing to integrate electrochemical sensors onto temperature-sensitive substrates such as OSTE+ polymers. Given that OSTE+ exhibits oxygen scavenging enabling oxygen at the microscale and mimicking…, here, electrochemical sensors were integrated to monitor dissolved oxygen in real-time. To this aim, gold and silver nanoparticle inks were inkjet-printed and sintered via photonic curing to form three-electrode electrochemical sensors directly on OSTE+. To validate the sintering approach, the electrochemical performance of thermally and photonically sintered electrodes was compared on polyethylene terephthalate (PET), a thermally stable substrate. The method was then applied to fabricate oxygen sensors directly on OSTE+ and integrated into a microfluidic device. Real-time dissolved oxygen levels were monitored for 18 h at three distinct locations within the microchannel, demonstrating the spatial resolution and temporal stability of the sensors. This low-temperature fabrication strategy overcomes the limitations of traditional high-temperature sintering techniques, expanding the range of materials suitable for sensor integration into MPS and OoC platforms.

## Materials and methods

### Fabrication of Teflon molds

Microfluidic devices were fabricated via soft lithography using custom-designed Teflon molds. The molds were designed using VCarve Pro 9.5 software (Vectric, UK) and were machined from 4 mm thick Teflon sheets using computer numerically controlled (CNC) milling system (Roland MDX-40A, Roland DG Corporation, USA). Three separate molds were fabricated, each corresponding to the three structural layers of the microfluidic devices: the upper layer (containing inlet and outlet ports), the middle layer (housing the microfluidic channel), and the bottom layer (serving as a sealing base with the integrated sensors). For toxicity assay, molds with cross-sectional dimensions of 4×12 mm and varying thicknesses of 1, 2, and 3 mm were used to fabricate the OSTE+ pieces.

### Device fabrication with oxygen-scavenging properties

Devices were fabricated using Ostemer® 322 Crystal Clear (Ostemer 322, Mercene Labs AB, Sweden). The polymer resin was prepared by mixing components A and B at a 1.09:1 weight ratio, followed by degassing in a vacuum chamber for at least 2h to eliminate bubbles. The degassed mixture was poured into the Teflon molds and excess resin was removed using a slab to achieve a smooth surface. The molds were then subjected to a two-step curing process. The first cure involved UV (365 nm) exposure at 1000 mJ/cm^2^ for 20 s. After the initial curing, the parts were demolded using flat tweezers and underwent a second thermal cure at either 30, 60, or 110 °C, depending on the experimental conditions. For control experiments, electrodes where integrated into cyclic olefin copolymer (COP, Microfluidic ChipShop, Germany) substrate, while 1 mm thick poly(methyl methacrylate) (PMMA) served as the structural layers for non-oxygen-scavenging microfluidic devices. The PMMA layers were laser-cut to match the dimensions of the OSTE+ microfluidic device. Both the oxygen-scavenging and control devices were assembled using double-sided pressure sensitive adhesive (ARCare 8939; Adhesive Research Europe). The layers where then aligned and bonded under 2 kN pressure for 10 min using a PW-H laboratory press (P-O Weber, Germany) at room temperature to ensure uniform adhesion.

### Inkjet printing

Printing was performed on UV-cured OSTE+ substrates to take advantage of the thiol-rich surface chemistry. Silver nanoparticle ink (DGP-40LT-15C; Advanced Nano Products, South Korea) was used for pseudo-reference electrode (pRE). Gold nanoparticle ink (Au-LT-20; Fraunhofer IKTS, Germany) was used for the working and counter electrodes (WE and CE). Both inks were deposited using the drop-on-demand Dimatix Materials Printer (DMP 2831; Fujifilm Dimatix; USA) featuring 16 nozzle with 10 pl drop size print head from which 1, 2, or 3 adjacent nozzles were selected for deposition. The drop spacing for both inks and the substrates used are as follows:

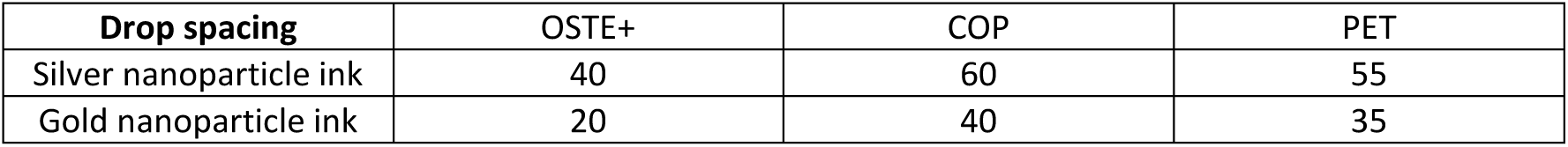

The printer system maintained the substrate at 30 °C during ink deposition for OSTE+. Comparative printing was performed on 125 µm thick polyethylene terephthalate (PET) and 1 mm COP substrates. Prior to printing, these substrates were exposed to oxygen plasma for 60 s at a power of 80 W using plasma treatment machine (Smart Plasma 80W-Oxygen, Plasma Technology, Germany) for improving ink adhesion. For PET and COP, the printer platen temperature was maintained at 60°C to promote solvent evaporation.

### Photonic and thermal sintering process

After ink deposition, sintering was carried out using a PulseForge Digital Thermal Processing™ system (NovaCentrix, USA). Samples were positioned 10 cm from the lamp and processed under ambient atmospheric conditions. Photonic curing parameters were optimized separately for silver and gold nanoparticle inks to minimize the number of pulses required for complete sintering. The pulse duration was fixed at 600 µs, while the applied voltage was varied from 250 V to 500 V, corresponding to an incident energy range of 0.163 to 2.53 J·cm⁻². For PET substrates, thermal sintering was performed in a convection oven at 150 °C for 1 hour. After curing, electrodes were passivated using a biocompatible, double-sided pressure-sensitive adhesive, ensuring that only the active electrode area remained exposed for subsequent measurements or functionalization. SimPulse software was used to simulate the thermal profile during photonic curing process.

### Pseudo-reference electrode development

All electrochemical measurements were conducted using a PalmSens4 potentiostat (PalmSens4 BV, Netherlands). The inkjet-printed silver electrodes were electrochemically chlorinated to stabilize their potential for use as pseudo-reference electrodes (pREs). For chlorination, all three electrodes (WE, CE, and pRE) were fully immersed in 0.1 M hydrochloric acid (HCl, Sigma-Aldrich). Cyclic voltammetry (CV) was then applied, scanning from −0.2 V to +0.2 V versus an external 2 mm in diameter Ag/AgCl reference electrode (DRIREF-2, World Precision Instruments), at a scan rate of 20 mV·s⁻¹, for a minimum of five cycles. To evaluate the electrochemical stability of the pRE, open circuit potential (OCP) measurements were performed. The pRE was immersed in 3 M potassium chloride (KCl, Sigma-Aldrich) solution, and the potential was recorded over a period of 1 hour. During OCP measurements, the pRE was connected as the WE, and an external Ag/AgCl reference electrode was used as the reference electrode to complete the electrochemical cell.

### Electrochemical characterization

Following electrochemical chlorination of the reference electrode, a 10 mM equimolar solution of potassium ferrocyanide and ferricyanide in 10 mM KCl was used to evaluate the performance of the printed electrode system. CV was performed over a potential window of −0.2 V to +0.5 V at a scan rate of 25 mV·s⁻¹. Three sequential configurations were assessed in the following order: first, the inkjet-printed WE was evaluated together with a commercial platinum wire CE and a commercial Ag/AgCl RE. Second, the platinum wire CE was replaced by the inkjet-printed CE. Finally, the fully printed three-electrode system was evaluated. The inkjet-printed CE was designed to be three times the size of the WE to avoid current limitation during measurements. In cases where the CV curve exhibited flattened redox peaks and reduced current response (indicative of limited electroactive surface area) an electrochemical activation step was applied. Activation consisted of five potential pulses alternating between −2.0 V and 0.0 V (versus Ag/AgCl), with 2s pulse durations over a total stimulation time of 16 seconds in PBS, based on a protocol from A. Moya et al[37]. Following activation, CV was repeated under identical conditions to assess improvement in the electrochemical response.

### Electrode calibration

Oxygen electrodes were calibrated by preparing solutions with varying oxygen concentrations, which were achieved by adjusting the sulfite concentrations in the electrolyte solution. The oxygen concentration in each solution was measured using an optical dissolved oxygen meter (HANNA Instruments) prior to electrochemical measurements to correlate the electrode signal with the known oxygen content. To investigate the oxygen reduction reaction, linear sweep voltammetry (LSV) was performed over a potential window from 0.2 V to −0.9 V, with a scan rate of 25 mV·s⁻¹. Following LSV, chronoamperometry was conducted at −0.8 V for each oxygen concentration. The potential was held for 60 s to allow the current to stabilize. The average current over the last 10 seconds of the chronoamperometric response was recorded and used to generate a calibration curve linking steady-state current to oxygen concentration, enabeling quantitative analysis of dissolved oxygen concentration.

### Microfluidic characterization of oxygen scavenging properties

Microfluidic experiments were conducted to characterize the oxygen scavenging properties of OSTE+. All experiments were performed at 25°C in an incubator, in 10 mM PBS solution. A custom-made PMMA holder was employed to securely seal the microfluidic tubing, preventing leakage and ensuring proper flow. Liquid perfusion was facilitated using a 4-channel peristaltic pump (Reglo ICC, ISMATEC, Cole-Parmer GmbH) with 2-stop pump tubing (Ismatec, PharMed® BPT) and silicone tubing (Freudenberg Medical). Microfluidic PEEK 1/16-inch fittings (XP-230X, IDEX Health & Science) were threaded into the holder, and stainless-steel tubing was used at the outlet to connect the microfluidic system to the oxygen sensor, preventing oxygen diffusion along the tubing. As a control oxygen concentrations at the outlet of the system were continuously measured using a commercial oxygen microsensor (UNISENSE, Denmark) integrated into a T-flow system and connected to a picometer controller (UNISENSE, Denmark). The microsensor was polarized at 0.8 V against an internal Ag/AgCl reference electrode. Before the experiments, the sensor was calibrated via a two-point procedure using an optical dissolved oxygen meter: one point in a solution at atmospheric oxygen concentration, and the second in an oxygen-free solution.

### Oxygen monitoring with inkjet-printed amperometric electrodes

Amperometric detection of dissolved oxygen was performed using a PalmSens4 potentiostat (PalmSens BV, Netherlands) in an incubator set at 25°C. Devices were fabricated with inkjet-printed electrodes embedded in OSTE+ microfluidic platforms and fabricated as described above. To evaluate the impact of thermal curing on oxygen scavenging and sensor performance, chips were post-cured for 1 hour at 30°C, 60°C, or 110°C prior to electrochemical testing. Measurements were performed directly after fabrication. A control device was fabricated with inkjet-printed electrodes on a COP substrate and PMMA microfluidic parts, which lacks oxygen scavenging capabilities, to evaluate the baseline oxygen profile. All devices were filled with 10 mM PBS, and chronoamperometry was performed by applying a constant potential of −0.8 V versus printed Ag/AgCl reference electrode. Current measurements were recorded every 5 minutes over a total duration of 14 hours.

### Oxygen consumption calculation

Oxygen consumption rate (mol O_2_/s) is directly proportional to the electrical current measured at the working electrode. To calculate oxygen consumption, the electrical current is first related to the number of electrons using the elementary charge constant (e = 1,6·10-19 C). From the stoichiometry of the reaction, which involves 4 electrodes to reduce O_2_ to H_2_O^-^, the number of oxygen molecules can be determined. Finally, using the Avogadro constant (NA = 6.022·1023 mol^-1^) the number of oxygen molecules is converted into moles of oxygen. This provides the direct oxygen consumption of an electrochemical sensor.

### Cytotoxicity assessment

To assess the cytotoxicity of Ostemer 322, two complementary experiments were performed. First, polymer samples were cured at 30 °C, 60 °C, or 110 °C to evaluate whether biocompatibility is preserved at temperatures lower than the manufacturer’s recommended 110 °C [38]. Second, to determine whether the amount of polymer influences biocompatibility and thereby the feasibility of fabricating devices of different dimensions under mild curing, additional samples of varying volumes (48, 96, and 144 mm³) were tested at 30 °C, since this condition favors thiol retention and oxygen scavenging. In both experiments, cured polymer pieces were incubated in MEM Alpha medium with GlutaMAX (Sigma-Aldrich), supplemented with 10% Fetal Bovine Serum (FBS, Sigma-Aldrich), at 37 °C in a humidified incubator for 24 h, with each condition prepared in triplicate. Media incubated under identical conditions without polymer exposure served as a negative control. Human intestinal colorectal adenocarcinoma cells (Caco-2, ECACC) were seeded in 24-well plates (Corning) and cultured for 5 days to reach confluence. Cells were then exposed to the conditioned media and incubated for 24 h at 37 °C in a humidified incubator. Cell viability was assessed using the MTT assay (3-(4,5-dimethylthiazolyl-2)-2,5-diphenyltetrazolium bromide, Corning). MTT reagent was added to each well at a concentration of 5 mg/mL and incubated for 4 h, after which 100 µL of DMSO was added to dissolve formazan crystals. Absorbance was measured at 570 nm using a UV-Vis miniature spectrometer (Ocean Optics) with a halogen light source (HL-2000, Ocean Optics), and cell viability was expressed relative to the untreated control.

### Data analysis

Data representation was performed using GraphPad Prism 10 (GraphPad Software). Statistical significance was assessed using unpaired t-tests; **** indicates p < 0.0001. Data are presented as mean ± standard deviation (SD). All graphs that represent real-time data were generated with an on-purpose Python-based developed program to automate the data analysis process. For the chronoamperometry data the values from the last 10 seconds were averaged and plotted.

## Results

### Low-temperature sensor integration concept

To enable integration of electrochemical sensors on OSTE+, a low-temperature fabrication strategy was developed. OSTE+ has a glass transition temperature of 69 °C, above which it softens and loses structural integrity, limiting high-temperature processing, such as thermal sintering of printed materials [26]. A three-electrode amperometric oxygen sensor was designed, consisting of a gold working electrode (WE), a gold counter electrode (CE), and a pseudo-reference electrode (pRE) based on Ag/AgCl. Oxygen reduction at the WE generates a current proportional to the oxygen concentration under an applied negative potential. This process can be observed by linear sweep voltammetry (LSV), which shows two distinct peaks at –0.4 and –0.8 V (LSV, **Figure 1a**). The first peak corresponds to the partial reduction of oxygen to hydrogen peroxide:

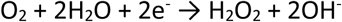

**Figure. 1.**
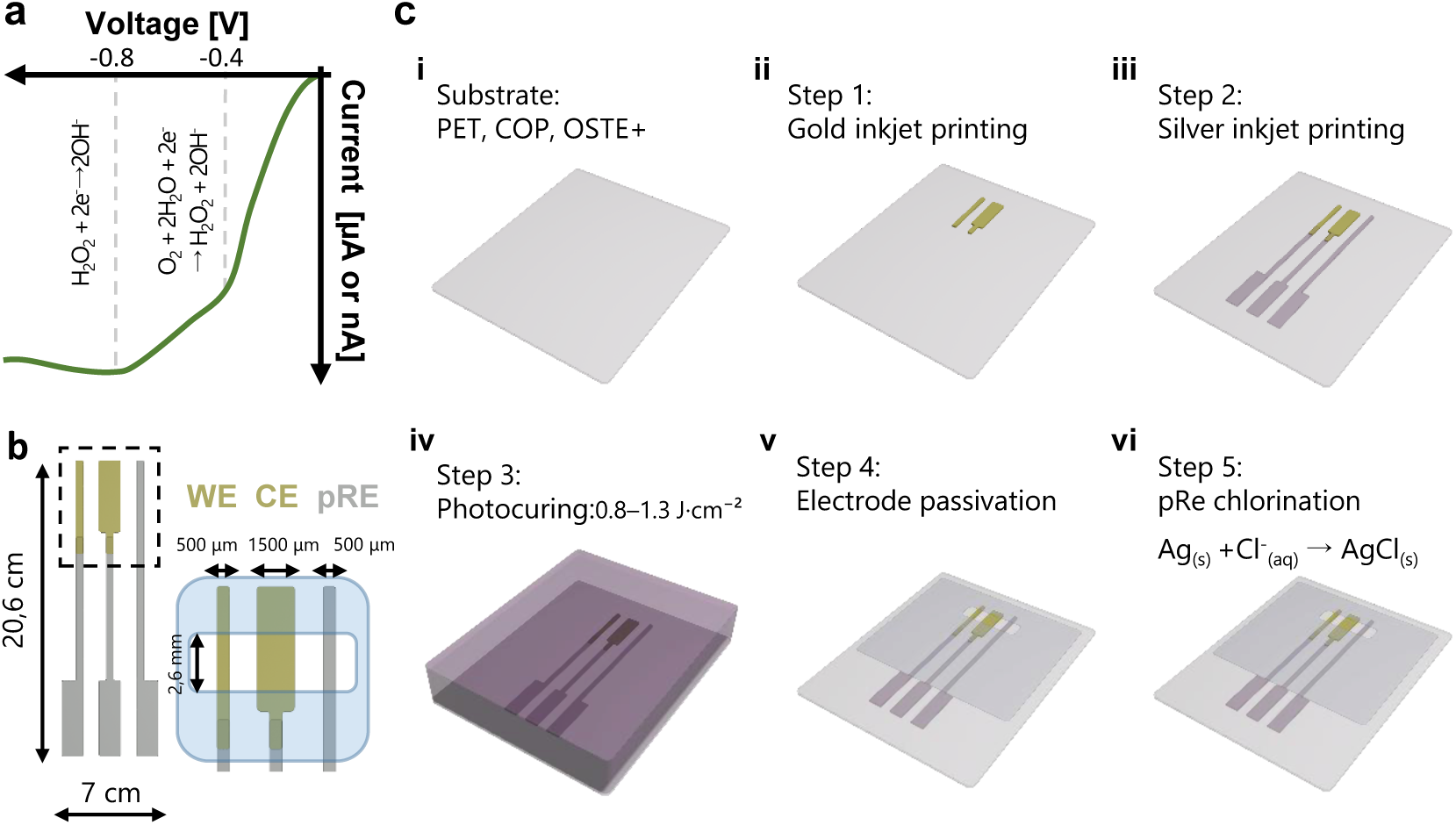
Oxygen sensor integration via inkjet printing and photonic curing for temperature-sensitive substrates. **a)** Schematic of a linear sweep voltammetry (LSV) displaying two reduction peaks at the WE when a negative voltage is applied. **b)** Illustration of an amperometric oxygen sensor with gold WE and CE electrode and an Ag/AgCl pRE. **c)** Fabrication steps: (i) Substrate selection, (ii) gold nanoparticle ink deposition, (iii) silver nanoparticle ink deposition, (iv) photonic curing for sintering, (v) passivation layer application to define active electrochemical areas, and (vi) pRE chlorination.

The second peak corresponds to the subsequent reduction of hydrogen peroxide to hydroxide ions:

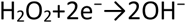

This redox reaction occurs in a basic environment, consistent with the pH of PBS (7.4). The electrodes were aligned in a single row, rather than in conventional circular designs, to minimize footprint and facilitate integration into space-limited microfluidic devices (**Figure 1b**). The WE and pRE each measured 500 µm in width, and the active sensing area was defined by a 2.6 mm opening, resulting in a WE area of 1.3 mm^2^. The CE was designed with three times the area of the WE to prevent current limitation.

The fabrication process is illustrated in **Figure 1c**. First (step i), substrate materials were selected, including OSTE+, polyethylene terephthalate (PET) and cyclic olefin polymer (COP). OSTE+ served as the primary substrate due to its unreacted thiol groups, which enable both strong metal adhesion and intrinsic oxygen-scavenging properties. PET was used to compare photonic and thermal sintering because it tolerates temperatures above 150 °C. COP was used as a non-reactive control to isolate substrate effects on oxygen dynamics. After substrate selection, gold and silver nanoparticle inks were deposited via maskless inkjet printing: gold ink was printed first to define the WE and CE (step ii), followed by silver ink for the pRE and interconnects (step iii). The printed inks were then sintered by photonic curing with a xenon lamp (300–1100 nm), enabeling rapid processing without thermal damage to the substrate (step iv). Optimized curing conditions are discussed later for each substrate. A pressure-sensitive adhesive was patterned to passivate non-electrochemical regions and define active sensing areas (step v). Finally, the silver pRE was electrochemically chlorinated to form a stable Ag/AgCl interface, ensuring a consistent reference potential, with measured OCP drift of 4.53 µV·h^-1^ for Ag and 1.21 µV·h^-1^ for Ag/AgCl (step vi, **SI Figure 1**). This protocol enables integration of functional electrochemical sensors on OSTE+ and other thermally sensitive materials.

### Photonic curing optimization

PET substrate was used for photonic curing optimization due to its compatibility with both thermal sintering (150°C for 1 hour) and pulse-light processing. Photonic sintering reduced the sheet resistance of gold and silver inks by 26% and 54%, respectively, compared to thermal curing (Figure 2a). This improvement arises from the high peak temperatures achieved during photonic curing, which can reach up to 400 °C at the film surface without significantly heating the underlying substrate (Figure 2b) [36]. This rapid, localized heating enables processing in milliseconds without compromising substrate integrity. However, steep thermal gradients generated internal stress in the ink films, leading to crack formation (Figure 2c). These effects are likely caused by rapid solvent evaporation and thermal stress triggered by high-intensity light pulses. To mitigate these effects, both curing energy (Figure 2d) and pulse number were optimized for both inks (SI Figure 2). Effective sintering required exceeding a minimum energy threshold to remove solvents, surfactants, and stabilizers, allowing nanoparticles coalescence. Optimal conductivity and minimal cracking occurred within an energy window of 0.8–1.3 J·cm⁻², while higher energies sharply increased crack density and ultimately disrupted electrical continuity (SI Figure 3). Pulse number also influenced performance, with two pulses yielding the lowest sheet resistance and variability. To compensate for surface defects and localized cracking, the number of printed layers was also optimized. Three layers for both gold and silver inks ensured sufficient conductivity while maintaining mechanical integrity (SI Figure 4). Morphological changes induced by curing were assessed by optical microscopy (Figure 2e). Prior to sintering, the films displayed a uniform texture. Thermal curing led to moderate thickening, reflecting nanoparticle coalescence. Under optimized photonic curing conditions, more pronounced changes in microstructure were observed, including localized particle fusion and surface texturing, particularly in silver films.

**Figure 2.**
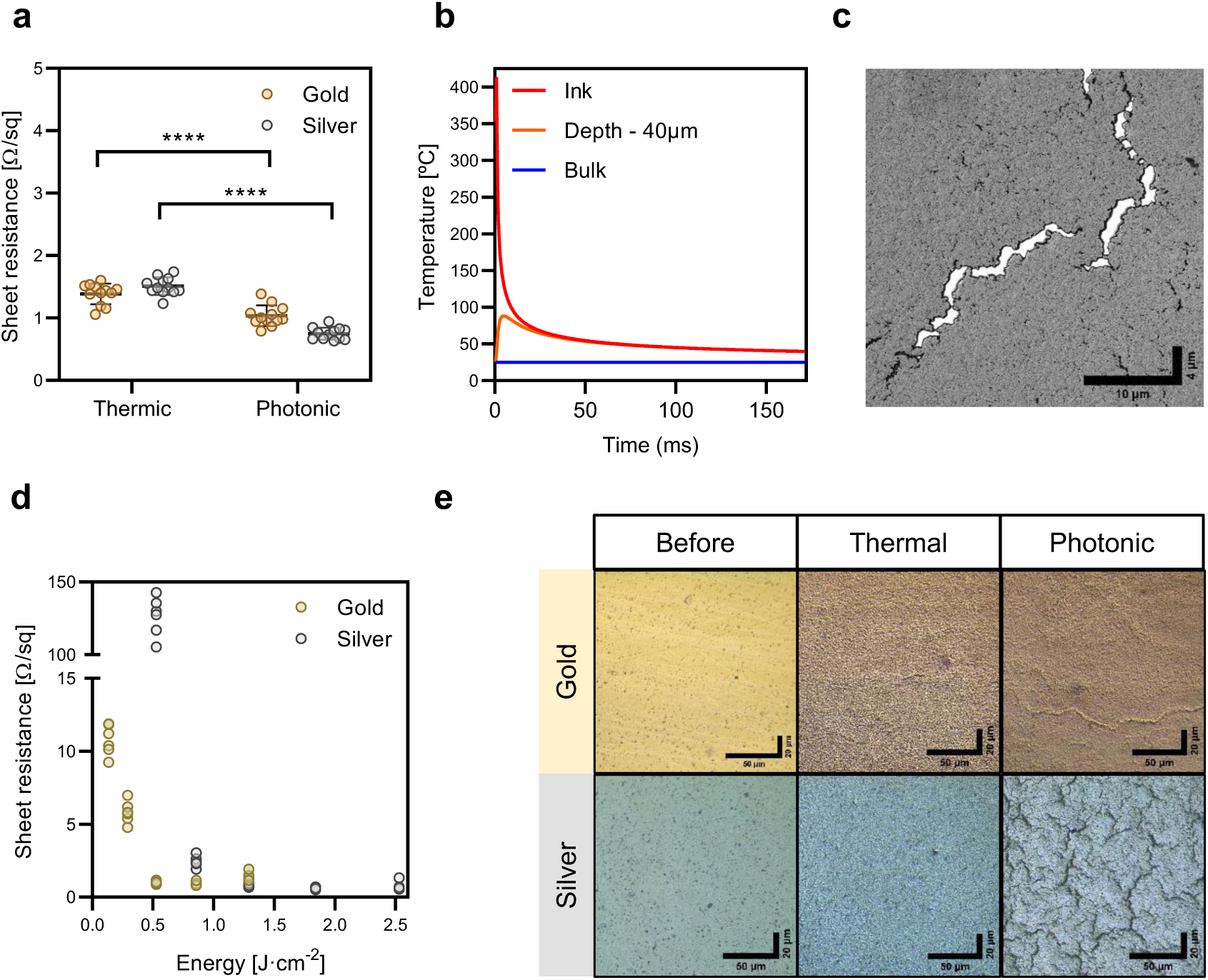
Evaluation of photonic curing parameters for inkjet-printed gold and silver inks on PET substrates. **a)** Sheet resistance values of inkjet-printed gold and silver on PET substrate after thermal curing and photonic curing. **b)** Simulated thermal profile during photonic curing (600 µs, 400 V, 1.3 J·cm⁻²) showing spatial temperature gradients across the film thickness. Peak temperatures occur at the film surface (red), while temperature rapidly decreases toward the substrate interface (40µm depth, orange), minimizing thermal load on the bulk material (blue). **c)** SEM image of a representative crack (∼1.5 µm wide) observed in gold ink after photonic curing. **d)** Sheet resistance values of gold and silver inks as a function of photonic curing energy. **e)** Microscopic images of inkjet-printed gold and silver ink: (left) before sintering, (middle) after thermal curing, and (right) after photonic curing.

**Figure 3.**
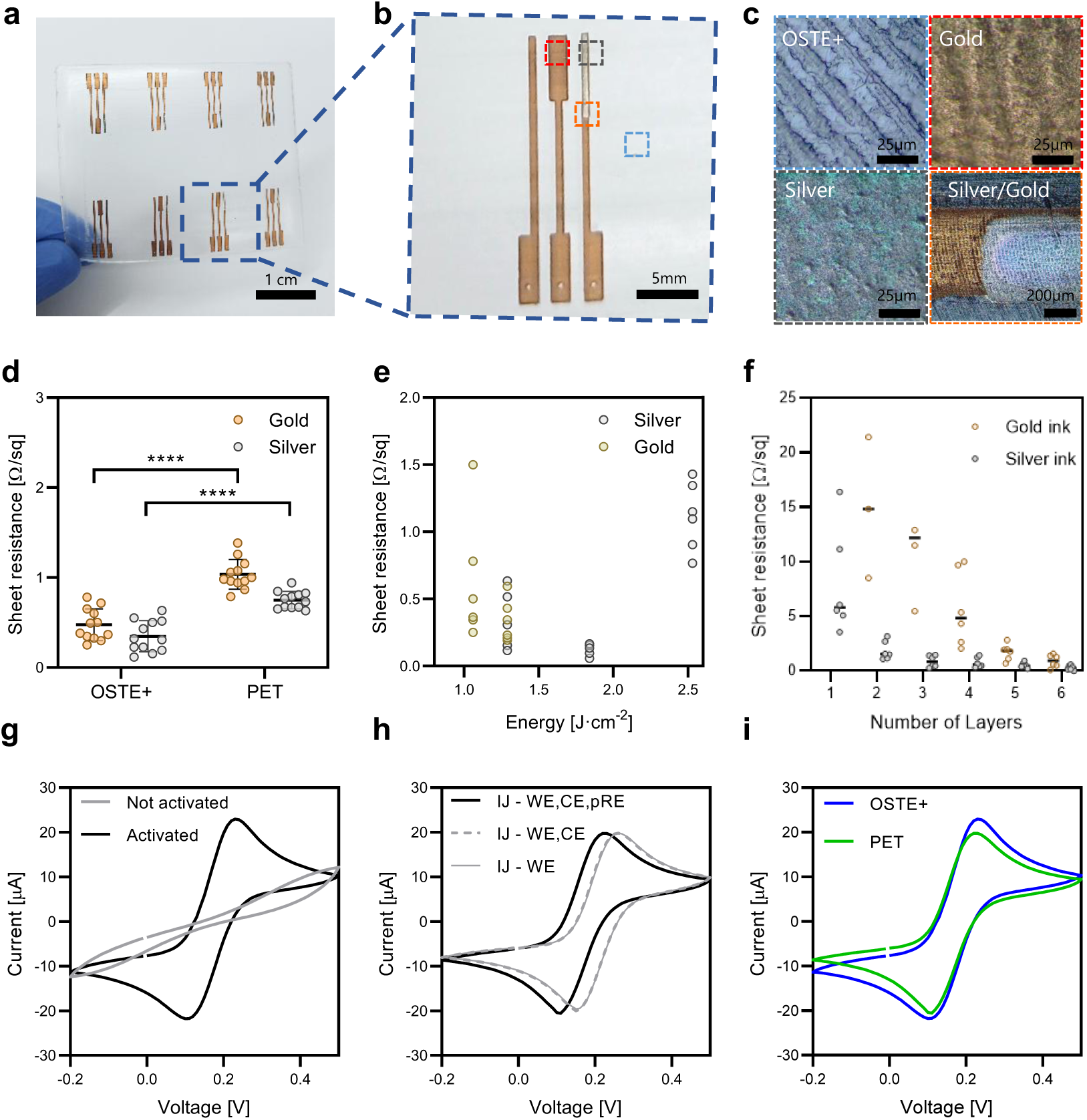
Inkjet-printed gold and silver electrodes on OSTE+. **a)** Photograph of a sensor array inkjet printed on OSTE+ substrate. **b)** Close-up of a single electrochemical sensor showing its three-electrode configuration: working electrode (WE), counter electrode (CE), and pseudo-reference electrode (pRE). **c)** Microscopy images showing: (top left) textured surface of cured OSTE+; (top right and bottom left) surface morphology of inkjet-printed gold and silver electrodes, respectively; and (bottom right) gold-silver interface. **d)** Comparison of electrode sheet resistance for gold and silver inks printed on PET and OSTE+ substrates after optimized photonic curing. **e)** Electrode sheet resistance as a function of photonic curing energy for gold and silver inks on OSTE+. **f)** Resistivity values of gold and silver inks as a function of the number of inkjet-printed layers on OSTE+ substrate. **g-i)** Electrochemical performance in 10 mM ferro/ferricyanide solution at a scan rate of 20mV·s^-1^ **g)** Cyclic voltammograms (CVs) before and after electrochemical activation of the WE in OSTE+. **h)** CVs comparing electrode configurations using inkjet-printed WE and commercial CE and pRE (line IJ-WE), using inkjet-printed WE and CE and commercial pRE (line IJ-WE,CE and all integrated inkjet-printed electrodes (line-WE,CE, and pRE) **i)** CVs comparing sensor performance on different substrates: OSTE+ and PET.

**Figure 4.**
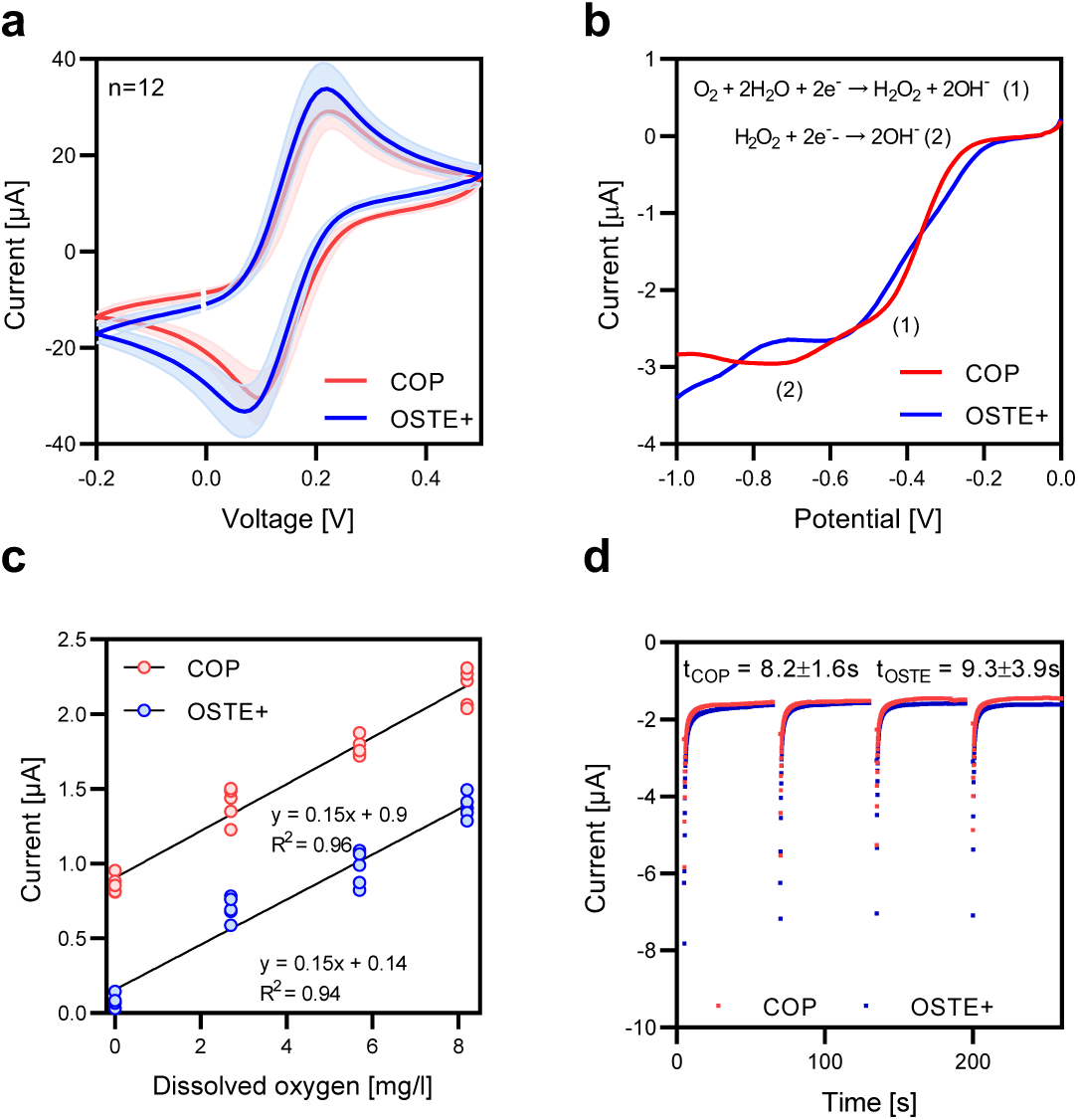
Electrochemical characterization and calibration for oxygen sensing. **a)** CVs of electrodes inkjet printed on OSTE+ and COP substrates (n=12). **b)** LSV on OSTE+ and COP in 10 mM PBS, showing the reduction of oxygen when a negative voltage window is applied from 0 to −1V. **c)** Calibration curves showing current density (µA·cm^-2^) response as a function of dissolved oxygen concentration (mg/l) for the same electrodes integrated into OSTE+ and COP substrates (n=4). **d)** Four consecutive chronoamperometric measurements in 10 mM PBS, each lasting 60 seconds, assessing the response time at a fixed dissolved oxygen concentration on OSTE+ and COP substrates (n=4).

### Electrode integration into OSTE+ polymer

OSTE+ retains unreacted thiol (-SH) groups after UV polymerization, which provide two key functionalities relevant for OoC/MPS systems. First, surface thiols allow direct covalent bonding with gold (Au) and silver (Ag) nanoparticles through gold-thiolate (Au–S) and silver-thiolate (Ag– S) interactions. This enables nanoparticle immobilization without additional surface modifications, providing an advantage over convetional substrates that typically require oxidative pretreatments such as oxygen plasma or chemical oxidants to improve adhesion [39]. Second, thiols confer oxygen scavenging capabilities, which are enhanced by lower thermal curing temperatures. This property allows precise tuning of oxygen concentration within the device microenvironement, enebeling the stablishement of physiologically relevant conditions [35]. Inkjet-printed electrochemical sensor arrays with a three-electrode configuration were fabricated directly on OSTE+ substrates (Figure 3a). Each unit included a WE, CE, pRE, and conductive tracks and pads (Figure 3b). The cured polymer’s textured surface, replicated from the Teflon mold (Figure 3c, top left), was preserved in the printed gold and silver electrodes (top right and bottom left), with a continuous and homogeneous Au–Ag junction (bottom right). Dimensional deviations of fabricated electrodes compared with the designed ones were below 5% (SI Figure 5). Electrodes on OSTE+ exhibited lower sheet resistance than those on PET, with average values reduced by 50% for both and gold silver inks (Figure 3d). This was achieved using a photonic curing energy of 1.29 J·cm^-2^ with six printed layers for gold and four for silver (Figure 3 **e and f**). However, the optimal curing energy window was narrower for OSTE+ (Figure 3e) than for PET (Figure 2d). This behavior may be linked to differences in substrate thickness and thermal properties. The greater thickness of OSTE+ (1 mm) compared to PET (125 µm) may lead to localized heat retention during curing, enhancing ink sintering but increasing the risk of thermal degradation. Cyclic voltammetry (CV) revealed passivated electrodes, characterized by low current response, which increased after electrochemical activation with well-defined redox peaks and a symmetric shape (Figure 3g). CV profiles were consistent across configurations: (i) printed WE with commercial CE and RE, (ii) printed WE and CE with commercial RE, and (iii) fully printed electrodes, showed consistent profiles (Figure 3h). A slight potential shift of 30 mV was observed with the printed pRE can be attributed to differences in chloride concentration. Finally, comparable CV responses on OSTE+ and PET confirmed equivalent electrochemical behavior using this low-temperature curing protocol regardless of the substrate (Figure 3i). The increased CV area observed for electrodes on OSTE+ compared to PET can be attributed to the textured surface of the cured OSTE+, which increases the effective electroactive surface area.

**Figure 5.**
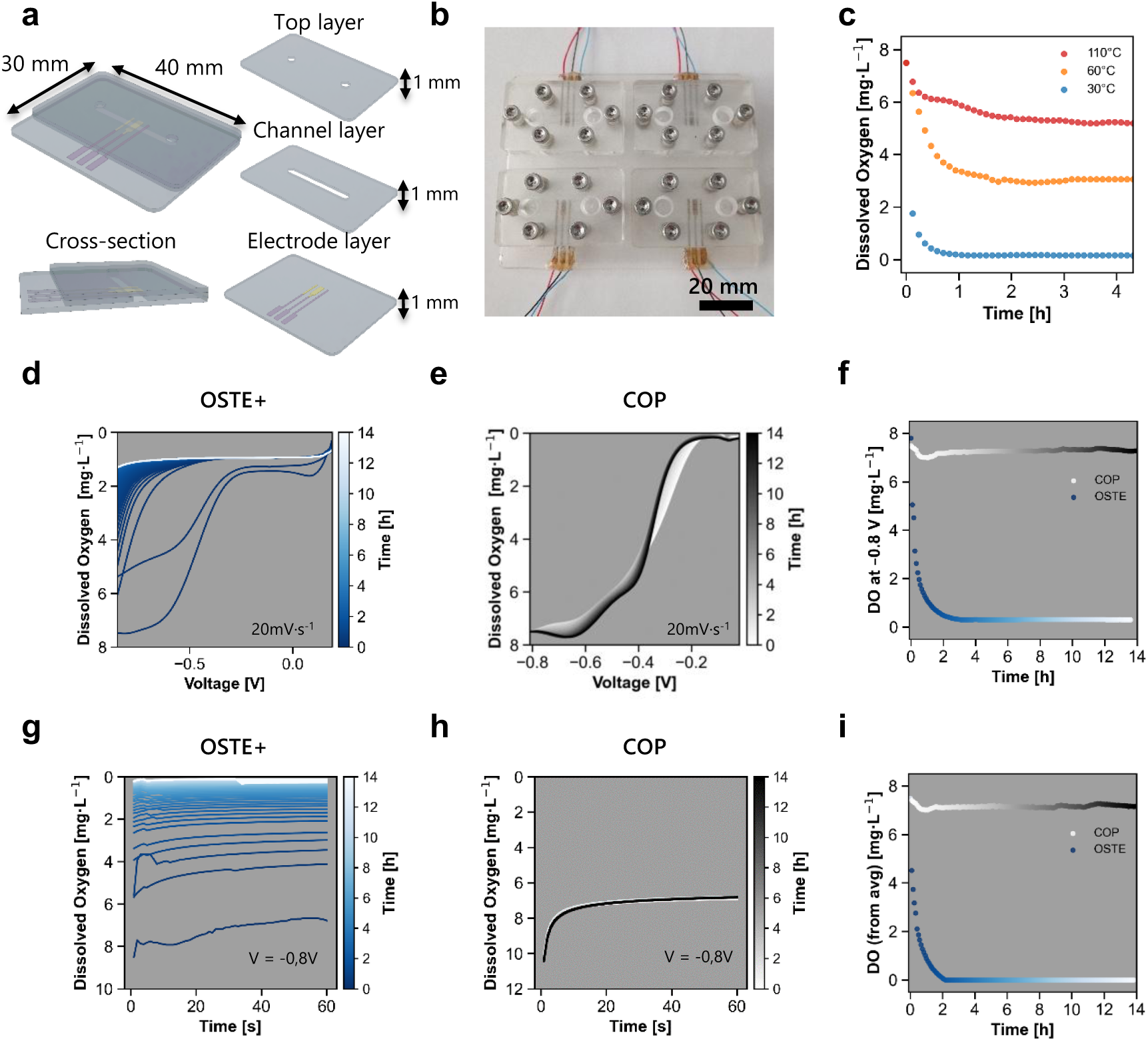
Oxygen reduction dynamics in OSTE+ versus COP microfluidic devices. **a)** Schematic of the device structure, comprising three layers: bottom (electrode layer), middle (microfluidic channel), and top sealing layer. A cross-sectional view depicts the internal channel structure. **b)** Photograph of four identical devices, each with a single integrated oxygen sensor, on one platform. **c)** Dissolved oxygen concentration over time measured in microfluidic devices cured at 30, 60, and 110°C during the second thermal curing step. **(d–e)** LSV in PBS from 0 to −0.8 V at 20 mV·s⁻¹, every 5 min over 14 h. OSTE+ devices (**d**) show progressive oxygen depletion, while COP devices (**e**) remain electrochemically stable, indicating no scavenging activity. **f)** Current at –0.8 V extracted from LSV, highlighting the difference in oxygen consumption between OSTE+ and COP. **g-h)** Chronoamperometry in PBS at –0.8 V over 60 s, every 5 min for 14 h. OSTE+ devices (g) show increasing oxygen reduction, while COP devices (**h**) remain stable. Devices fabricated with **(g)** OSTE+ again exhibit oxygen reduction, while **(e)** COP shows stable response. **i)** Comparison of OSTE+ and COP dynamics based on the average current during the final 10 s of each chronoamperometric measurement.

### Electrochemical electrode characterization and calibration

Following electrode integration into OSTE+ for dissolved oxygen measurements, the same fabrication protocol was also applied to COP, a polymer lacking intrinsic oxygen scavenging properties (SI Figure 6). COP served as a control to isolate the contribution of OSTE+ thiol chemistry to oxygen reduction. Moreover, since electrochemical sensing consumes dissolved oxygen, using COP confirmed that the oxygen depletion observed on OSTE+ originates from its thiol-based surface chemistry rather than electrochemical consumption. CVs from electrodes on both substrates exhibited similar redox peak shapes and currents, indicating reproducible and electrochemically equivalent sensor performance (n = 12, Figure 5a). To qualitatively confirm oxygen reduction at the WE, LSV was conducted on both substrates (Figure 5b), revealing a current increase with negative potential consistent with dissolved oxygen reduction. Calibration curves exhibited a linear relationship between current response and dissolved oxygen concentration (0–8 mg/L, equivalent to 0–21% atmospheric O₂), with a sensitivity of 0.15 μA·l·mg^-1^ for both substrates. (Figure 5c). A slight offset toward more negative potentials was observed for COP, suggesting less favorable oxygen reduction kinetics compared to OSTE+. Sensor response time was evaluated through four consecutive chronoamperometric measurements at constant oxygen concentration. Devices on OSTE+ showed an average response time of 9.3 ± 3.9 s, while those on COP responded with 8.2 ± 1.6 s (Figure 5d). These data demonstrate that both substrates support reliable low-temperature sensor integration with equivalent sensitivity and response times. The favorable thiol chemistry in OSTE+ likely facilitates strong metal–thiol interactions, which may contribute to enhanced oxygen reduction kinetics.

**Figure 6.**
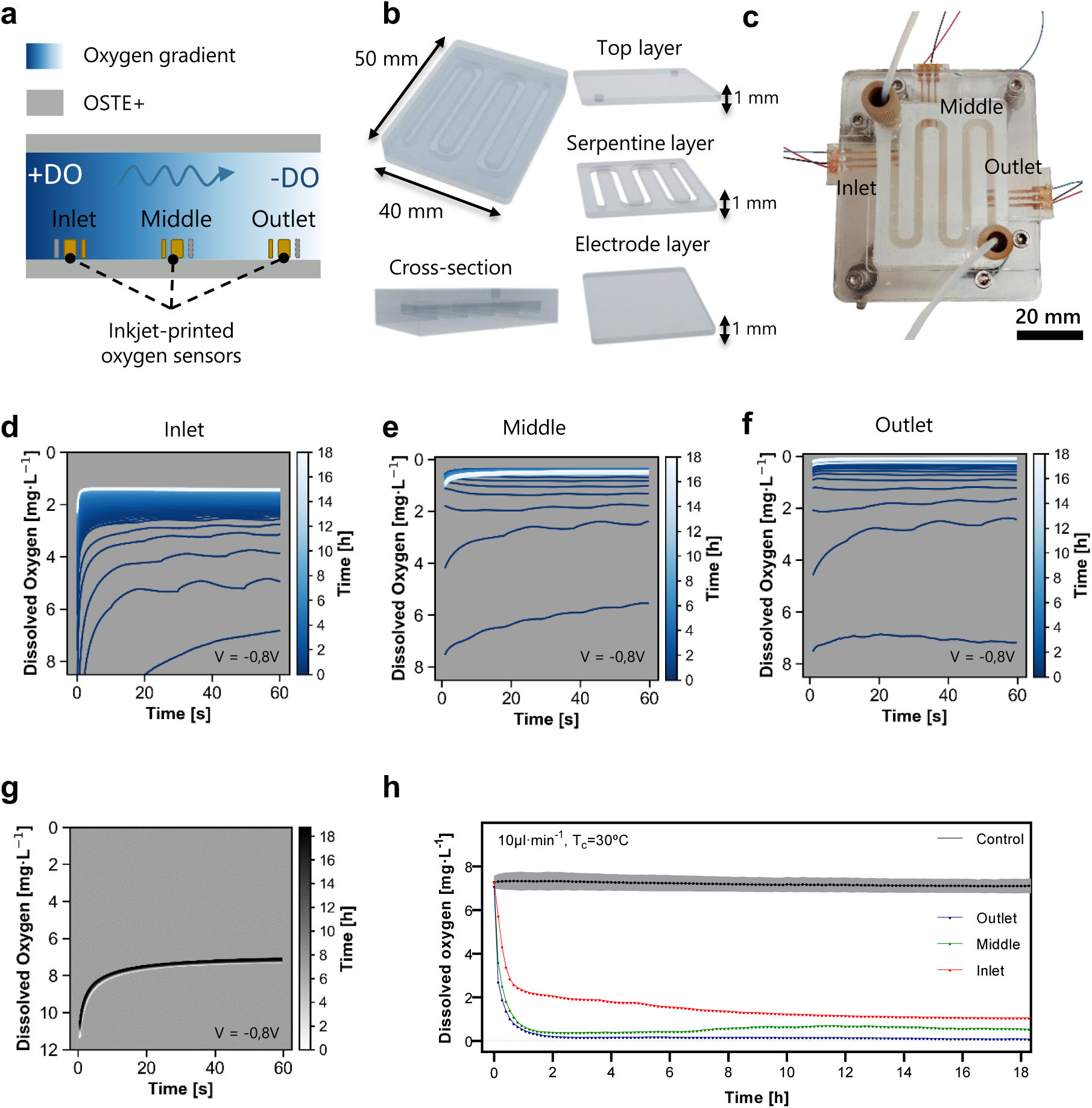
Integration of electrochemical oxygen sensors into an OSTE+ microfluidic device for spatial and temporal monitoring under flow. **a)** Schematic depicting spatial oxygen gradients along the channel, with electrochemical sensors positioned at the inlet, middle, and outlet for spatially resolved sensing. **b)** Photograph of the assembled microfluidic device fabricated from OTE+. **c)** Schematic of the device structure, comprising three layers: bottom (electrode layer), middle (microfluidic channel), and top sealing layer. A cross-sectional view depicts the internal channel structure. **d–f)** Chronoamperometric oxygen measurements acquired over 60 seconds every 5 min over 18h at the inlet **(d)**, middle **(e)**, and outlet **(f)** electrodes of the OSTE+ device. Current signals were converted to oxygen concentrations using calibration curves. **g)** Chronoamperometric response from a COP-based control device under equivalent conditions. **h)** Long-term oxygen measurements over 18 hours at the inlet, middle, and outlet electrodes under continuous flow (10 µL·min⁻¹) and a thermal curing temperature of 30°C. A COP-based device was included as a non-scavenging control (n=3).

### Comparison of oxygen scavenging efficiency in OSTE+ and COP

To evaluate a room-temperature sensor integration strategy into OSTE+ polymer, a multilayered microfluidic device with an integrated inkjet-printed oxygen sensor was fabricated. The device consisted of three casted OSTE+ layers, with a channel (width: 2.6 mm, length: 22.6 mm, height: 1 mm) incorporated in the middle layer, providing a total oxygen-scavenging surface area of 58.8 mm² (Figure 5a). The sensor was positioned at the center of the channel to enable real-time monitoring of dissolved oxygen. For fabrication and testing, four identical units were prepared in parallel on a single platform (Figure 5b). Devices cured at 30°C demonstrated the highest oxygen-scavenging efficiency, consistent with maximal retention of unreacted thiol groups. Increasing the curing temperature to 60°C and 110°C resulted in progressively reduced scavenging capacity (Figure 5c). LSV scans from 0 to –0.8 V in PBS were performed every 5 min over 14 hours under static conditions. OSTE+ devices with curing temperature of 30°C showed a progressive decrease in dissolved oxygen, indicating spontaneous oxygen scavenging upon contact with aqueous media (Figure 5d). In contrast, COP controls exhibited stable LSV profiles with two well-defined reduction peaks (Figure 5e). Current quantification at –0.8 V, corresponding to complete oxygen reduction, revealed rapid oxygen depletion within 2 hours for OSTE+, while COP showed no significant change (Figure 5f). Chronoamperometry at –0.8 V for 60 s every 5 min confirmed these trends (Figure 5g**,h**), with dissolved oxygen averaged during the last 10 s further validating oxygen depletion in OSTE+ and stability in COP (Figure 5i).

Cytotoxicity assays using Caco-2 cells exposed to polymer supernatants from samples cured at 30, 60, and 110 °C revealed no significant reduction in viability compared with untreated controls, confirming that OSTE+ remains biocompatible even at curing temperatures lower than the manufacturer’s recommended 110 °C (**SI** Figure 7a). To further examine whether polymer amount influences cytotoxicity, additional assays performed at 30 °C with varying volumes (48, 96, and 144 mm³) also showed no detectable effect on cell viability (**SI** Figure 7b). Together, these results confirm the overall biocompatibility of OSTE+ and demonstrate that oxygen scavenging capacity is modulated by curing temperature, reflecting differences in thiol retention. This tunable behavior is particularly important for generating physiologically relevant oxygen profiles levels in MPS and OoC platforms.

### Real-time monitoring of oxygen gradients in an OSTE+ microfluidic device for OoC models

To resolve spatial oxygen dynamics under perfusion, a multilayer microfluidic device with three integrated inkjet-printed sensors was fabricated (Figure 6a). All layers were cast from OSTE+ and measure 1 mm in thickness. The middle layer contains a serpentine channel (width: 2.6 mm, length: 205 mm), providing 550 mm² of oxygen-scavenging surface area (Figure 6b). The bottom layer incorporates three sensors at the inlet, middle, and outlet positions, while the top payer serves as a cover (Figure 6c). Initial characterization of oxygen-scavenging performance employed a commercial electrochemical sensor at the outlet under flow. Evaluation of curing temperature, flow rate, channel surface area, and post-fabrication time revealed distinct dissolved oxygen modulation profiles (**SI** Figure 8). Subsequent experiments with integrated sensors were performed on devices cured at 30°C with a 550 mm² surface area and immediately after fabrication to enhance oxygen scavenging and operated at a flow rate of 10 µl·min⁻¹, optimizing residence time for oxygen scavenging. Chronoamperometric measurements at inlet, middle, and outlet electrodes over 18 hours (Figures 6d**–f**) showed rapid oxygen depletion, establishing a stable gradient within the first hour (Figure 6h). Dissolved oxygen decreased from 7.5 mg·L⁻¹ to ∼1.0, 0.5, and <0.08 mg·L⁻¹ at the inlet, middle, and outlet, respectively. These dissolved oxygen levels at the outlet are in agreement with measurements obtained using the commercial sensor. COP-based controls maintained stable oxygen levels (∼7.5 mg·L⁻¹) throughout the experimental time, confirming that scavenging is specific to OSTE+. These results demonstrate the capability to precisely control and monitor physiologically relevant oxygen gradients with OSTE+ polymers, highlighting its potential for advanced OoC/MSP applications requiring oxygen control.

## Conclusion

This study demonstrates a low-temperature photonic curing protocol for the direct integration of inkjet-printed electrochemical sensors within OSTE+ microfluidic devices. OSTE+ is a thiol-rich polymer that provides unique advantages for sensor integration. Its residual thiol groups enable direct immobilization of conductive nanoparticles such as gold and silver via strong thiol-metal interactions, facilitating stable electrode fabrication. The optimized curing energy of approximately 1.29 J·cm⁻² achieved minimal electrode resistivity and cracking, producing high-fidelity electrode patterns with reproducible electrochemical performance. Notably, electrodes on OSTE+ exhibited approximately 50% higher conductivity compared to those on conventional PET substrates, without requiring any substrate pretreatment. Moreover, the protocol was successfully applied across multiple substrates, including PET and COP all relevant for OoC and MPS.

By combining the intrinsic oxygen-tunable capacity of OSTE+ with low-temperature sensor integration, this work presents a strategy for both recreating and monitoring precise oxygen levels on-chip. This capability is particularly critical for modeling organs with specific oxygen tensions (e.g., gut, liver, brain), pathophysiological microenvironments (e.g., tumors, ischemic regions such as stroke), or creating controlled hypoxic and near-anoxic conditions in vitro. The fabricated OSTE+ microfluidic devices enable real-time and on-chip monitoring of dissolved oxygen levels, while simultaneously modulating oxygen concentrations, representing a significant advance towards material-driven, multifunctional platforms for advanced OoC and MPS.

## Acknowledgements

The authors acknowledge the support from the Ministerio de Ciencia Innovación y Universidades (MICIU/FEDER, EU), in Spain, through GUMICHIP project (RTI2018-096786-B-I00). Denise Marrero acknowledges that this work has been done in the framework of the Ph.D. program in Electrical and Telecommunication Engineering at the Universitat Autònoma de Barcelona and was supported by the FPI Ph.D. fellowship (PRE2019-089214). This work has made use of the Spanish ICTS Network MICRONANOFABS partially supported by MICINN and the ICTS ‘NANBIOSIS’, more specifically by the Micro-NanoTechnology Unit of the CIBER in Bioengineering, Biomaterials and Nanomedicine (CIBER-BBN) at the IMBCNM. Cytotoxicity experiments were performed at Servei de Cultius Cel·lulars, Producció d’Anticossos i Citometria (SCAC) belonging to Universitat Autònoma de Barcelona (UAB). This work was also supported by Generalitat de Catalunya (2021SGR00495).

## Supplementary Information

**SI Figure 1.**
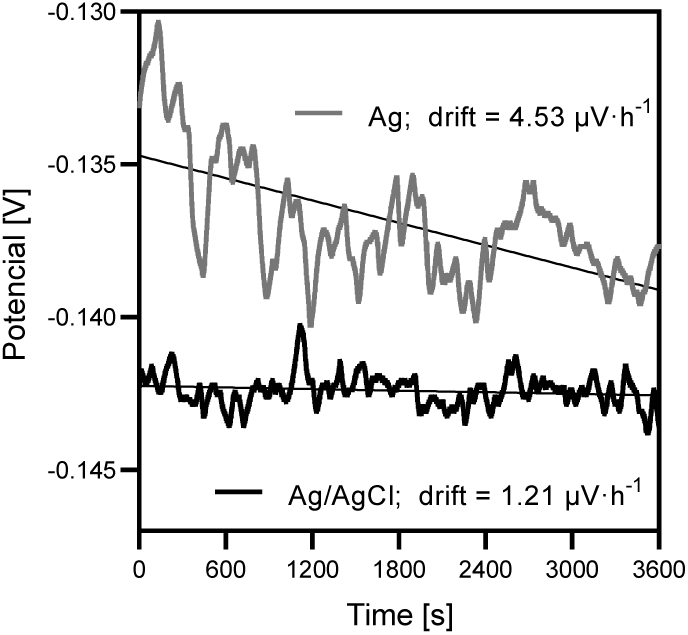
pRE chlorination. Open circuit potential measurements of the reference electrode with bare silver and developed pseudo-reference electrode with Ag/AgCl.

**SI Figure 2.**
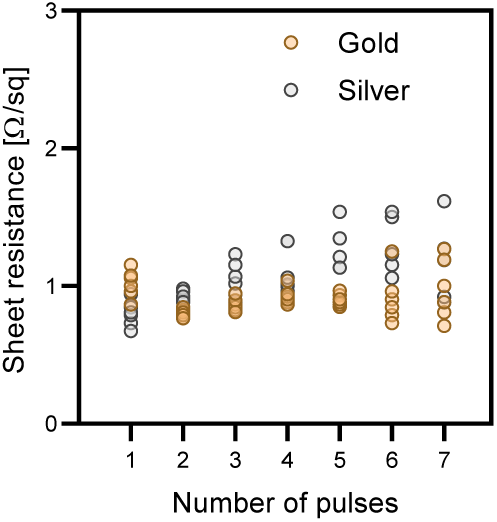
Optimization of light pulses. Sheet resistance of gold and silver inks as a function of the number of pulses in PET substrate.

**SI Figure 3.**
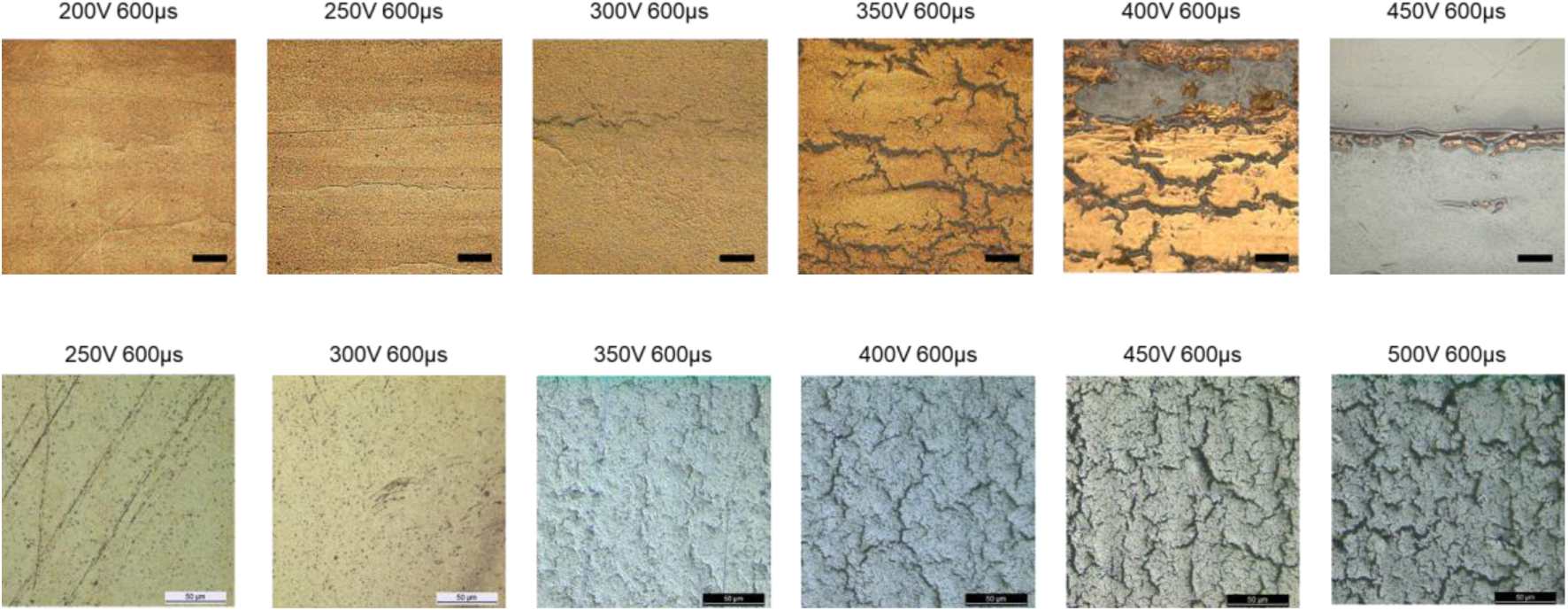
Optimization of photocuring energy. Representative optical microscopy images of inkjet-printed gold (top) and silver (bottom) on PET, illustrating the effect of increasing photocuring energy on crack formation. Higher photocuring energy promotes crack formation in both inks, with silver requiring a higher energy threshold than gold. Crack formation is observed at 300 V for gold ink and 350 V for silver ink.

**SI Figure 4.**
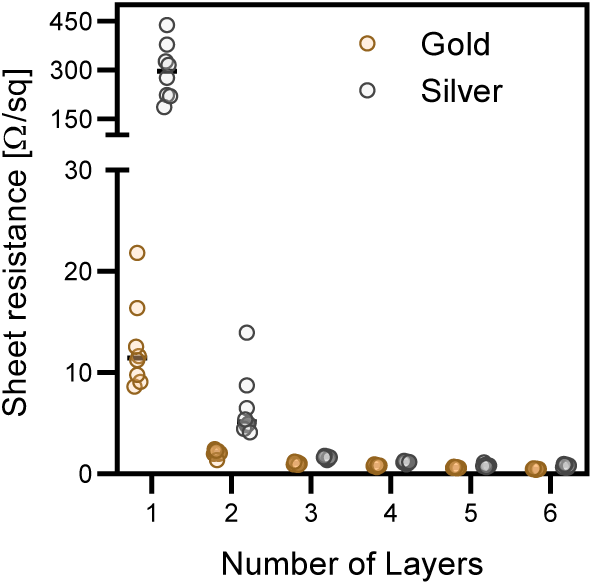
Optimization of ink layers. Sheet resistance of gold and silver inks as a function of the number of inkjet-printed layers on PET substrate.

**SI Figure 5.**
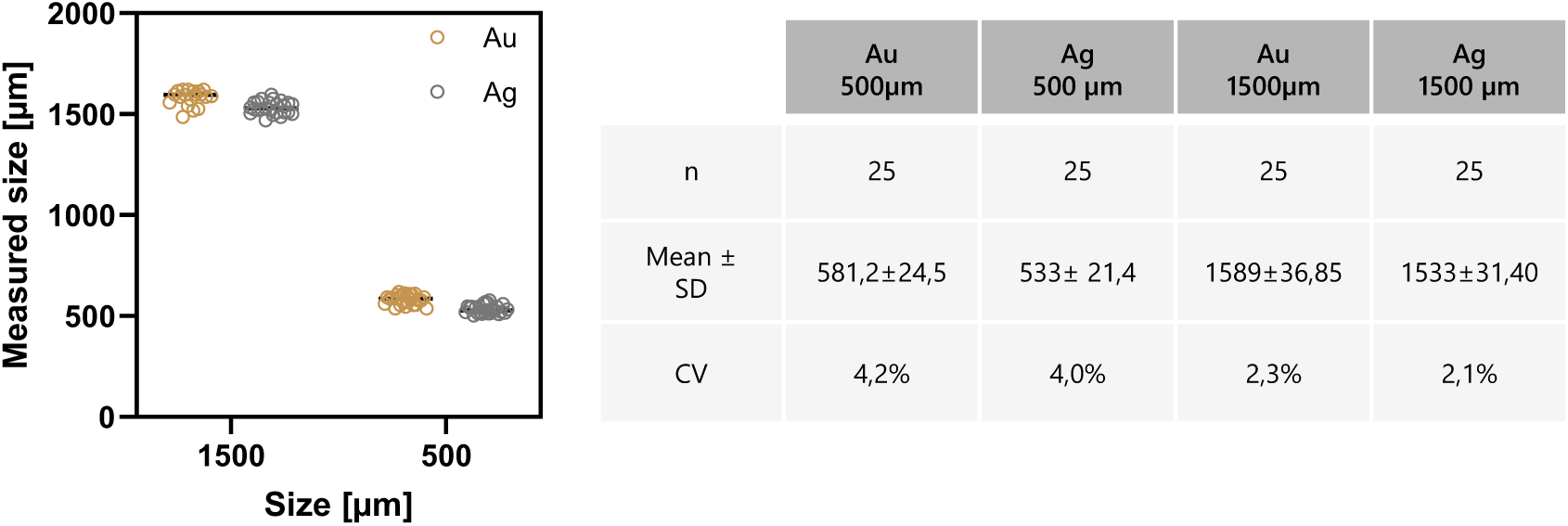
Geometrical accuracy of inkjet-printed electrodes on OSTE+. **a)** Comparison between the designed electrode dimensions (1500 µm and 500 µm) and the actual printed sizes measured by optical microscopy. **b)** Table summarizing mean ± standard deviation (SD) and coefficient of variation (CV) for measured electrode sizes (n=25 per geometry).

**SI Figure 6.**
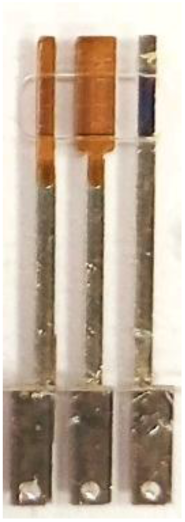
Inkjet-printed gold and silver electrodes on COP.

**SI Figure 7.**
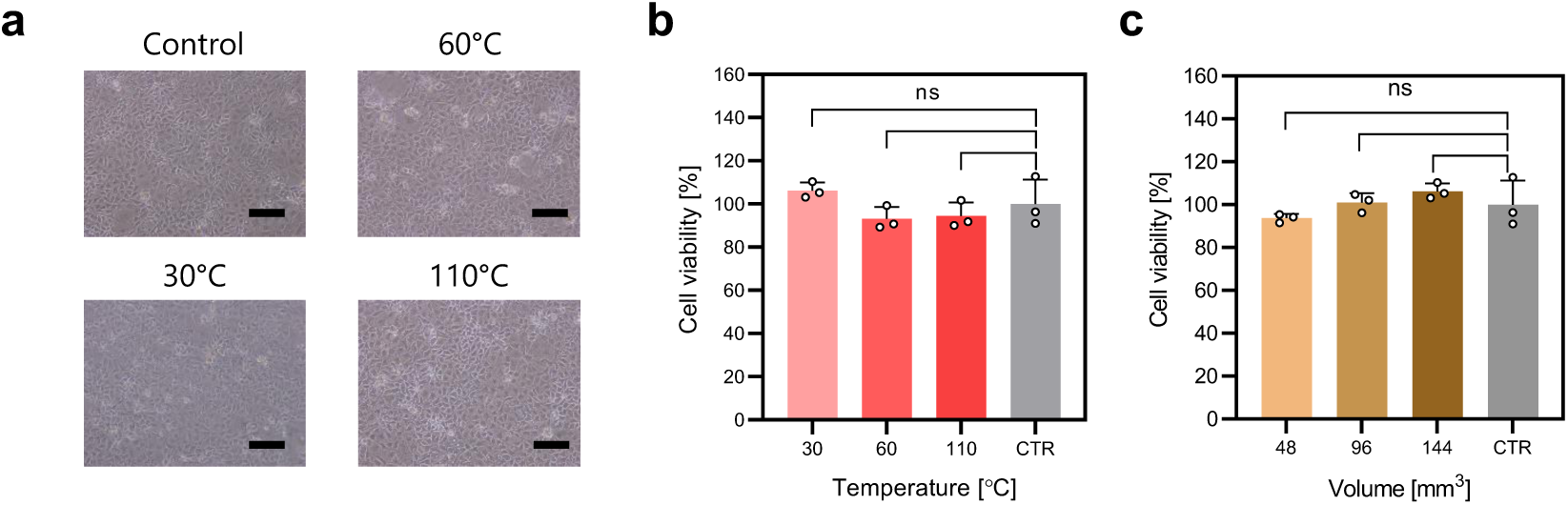
Cytotoxicity assessment test with Caco2 cells. **a)** Representative optical microscopy images of Caco2 cells cultured with Ostemer 322 Crystal Clear supernatant cured at different temperatures (scale bar = 100 µm). Cell viability of Caco2 cells exposed to medium in contact with Ostemer 322 Crystal Clear pieces cured at **b)** different temperatures (110, 60, and 30 °C) and of **c)** varying sizes (48, 96, 144 mm^3^) curet at 30 °C for 24 hours.

**SI Figure 8.**
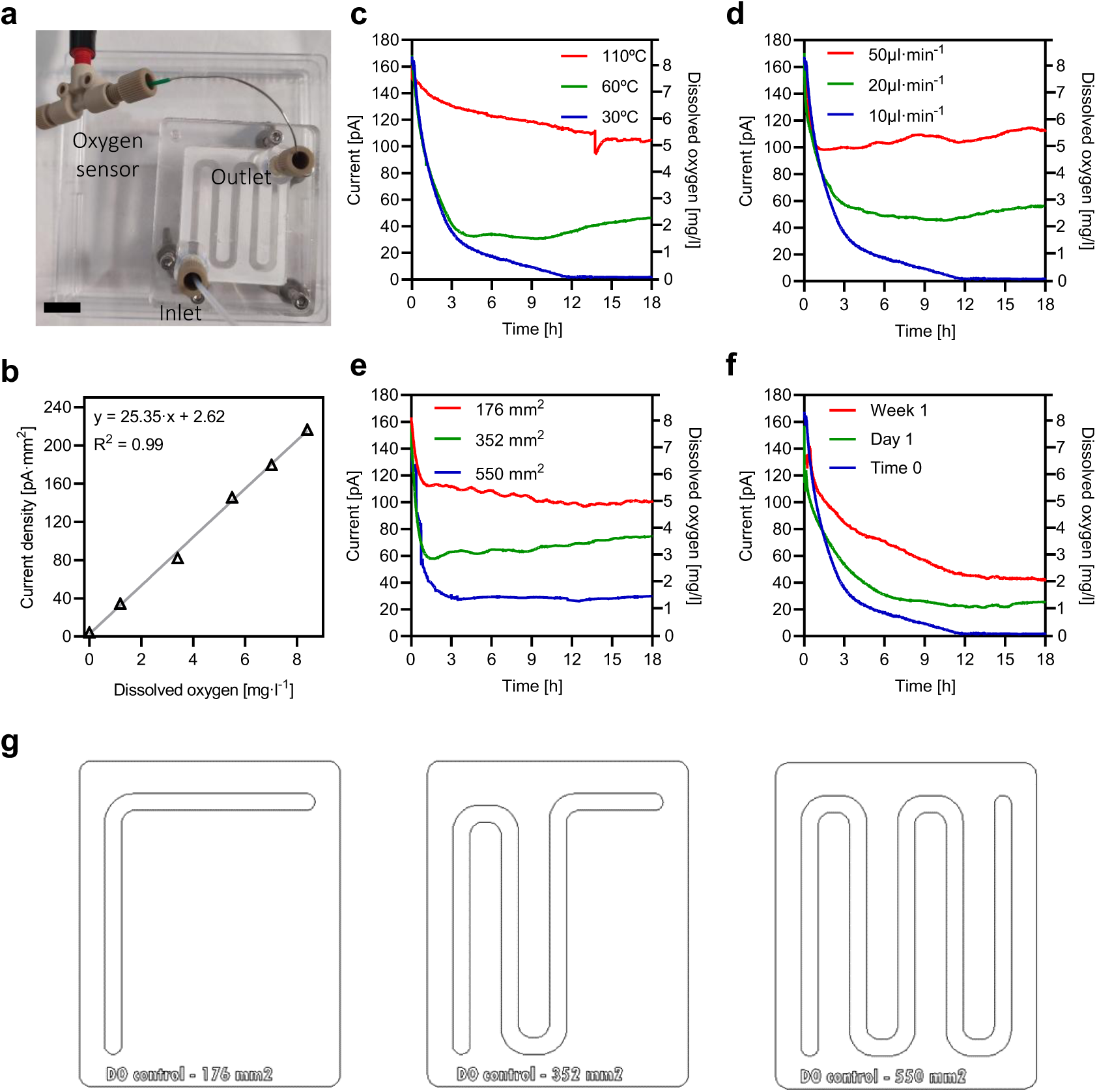
Oxygen levels characterization under flow conditions using a commercial sensor. a) Experimental microfluidic device setup to generate varying oxygen levels under flow conditions. Oxygen levels are measured using a commercial oxygen sensor at the output of the microfluidic device (scale bar = 10 cm). b) Unisense oxygen microsensor calibration curve. Oxygen levels are modulated by adjusting c) temperature d) flow rate, e) surface area, f) time after fabrication. g) Device designs with varying channel geometries to modulate oxygen-scavenging efficiency, corresponding to the results in panel e).

